# Superspreaders increase deleterious mutant burden and can accelerate the evolution of complex traits in pathogens

**DOI:** 10.64898/2026.04.24.720758

**Authors:** Samrat S. Mondal, Anastasia E. Madsen, Dominik Wodarz, Natalia L. Komarova

## Abstract

Degree heterogeneity in contact networks is known to accelerate the spread of infectious diseases through the presence of superspreaders, but its evolutionary consequences remain less understood. Here we study how network heterogeneity shapes the fate of competing pathogen strains in a stochastic susceptible–infected–susceptible framework. We show that heterogeneous networks act as strong suppressors of selection: both advantageous and disadvantageous mutants exhibit fixation probabilities close to neutral expectations, in stark contrast to well-mixed populations. We derive an analytical theory that captures this effect through a single suppression factor determined by network structure and infection dynamics, and validate it against simulations on synthetic and empirical contact networks. Mechanistically, suppression arises because most transmission events are effectively neutral, while selection acts only in rare configurations. As a consequence, heterogeneous networks substantially increase the persistence of deleterious mutants and elevate mutation–selection balance, but they can either accelerate or decelerate multi-step evolutionary processes such as fitness valley crossing. Our results reveal a fundamental trade-off induced by superspreaders: While they enhance epidemic spread, they weaken selective pressures and thereby promote evolutionary diversification.

---

Evolutionary theory on graphs has emerged as an area of intense interest, because network population structures better represent interactions in human, animal, and cellular populations compared to traditional well-mixed models [1, 2]. Network structure can amplify or suppress selection depending on characteristics like the heterogeneity of connections [1, 2]. It is especially relevant to infectious disease transmission, where highly connected individuals, called super-spreaders [3–5], can disproportionately drive transmission and substantially increase the basic reproductive number of a pathogen [3]. Previous numerical work has shown, however, that super-spreaders can paradoxically slow the emergence of advantageous mutants [6]. An analytical understanding of how network structure and super-spreaders jointly shape evolutionary dynamics is, however, currently lacking. In particular, no analysis exists for the evolution of deleterious mutants, which matter for future adaptation under environmental change and for adaptation involving complex traits that require multiple mutational hits. In this letter we provide comprehensive results for such evolutionary processes on networks with super-spreaders, filling this gap.

We consider susceptible–infected–susceptible dynamics [7] on a symmetric weighted host network *A* of size *N*, where *A*_*yx*_ denotes the weight of the connection between nodes *y* and *x*. Each node can be either *infected* or *susceptible*. We assume the existence of two variants of the pathogen, *wild-type* and *mutant*. Depending on its infection type, an infected node *y* is characterized by the infectivity *β*_*y*_ ∈ {*β*_w_, *β*_m_}, where *β*_w_ = *β* and *β*_m_ = (1 + *s*)*β* denote the infectivity of the wild-type and mutant, respectively. The parameter *s* specifies the selective advantage: *s >* 0, *s* = 0, and *s <* 0 indicate advantageous, neutral, and deleterious mutants, respectively. An infected node *y* attempts infection at rate *β*_*y*_*k*_*y*_ by choosing a neighbor *x* with probability *A*_*yx*_*/k*_*y*_, where *k*_*y*_ = ∑_*z*_ *A*_*yz*_. If *x* is susceptible, it becomes infected; otherwise, the attempt terminates. Infected nodes recover at rate *γ*.

The ratio *β/γ* controls basic reproductive number *R*_0_ [8], thereby determining infection persistence on the network. In all simulations, the infectivity is chosen sufficiently large for the wild type to maintain a nonzero steady-state with average infected population size *N*_i_. Simulations begin with few wild-type infections; after reaching steady state, a single mutant is introduced at a randomly chosen infected node. The mutant fixation probability *ρ*(*s*) is the probability that the mutant eventually takes over the infected subpopulation (Appendix A).

We first examine how network structure affects evolution through the fixation probability of *single-hit* mutants and ask whether selection is altered. Consider two synthetic degree-heterogeneous networks: A star graph [1], consisting of one central node connected to multiple leaf nodes, each having the central node as its only neighbor, and a Barabási–Albert scale-free network [9]. To determine whether selection strength is changed by the network structure, in FIG. 1(a-d) we plot the probability of mutant fixation on these two networks (blue dots) and compare it to that for a fully connected network [10] (black lines). We observe both networks strongly suppress selection for advantageous and disadvantageous mutants, bringing the fixation probabilities close to the neutral expectation, ∼ 1*/N*_i_ [11] (gray lines).

**FIG. 1.**
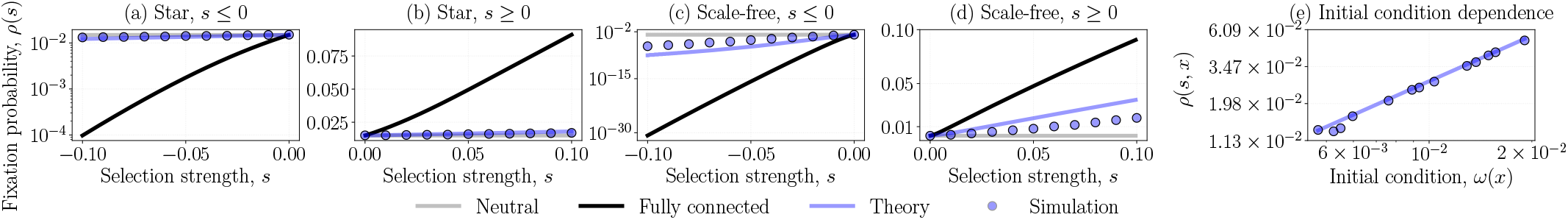
Heterogeneous networks strongly suppress selection. In panels (a)–(d), we plot the fixation probability as a function of selection strength for a star graph with *N* = 10^2^ (a,b) and a scale-free graph with *N* = 10^3^ (c,d), separating the results for deleterious (a,c) and advantageous mutants (b,d). In panel (e), we show the initial-condition-dependent fixation probability *ρ*(*s, x*) on the scale-free graph for a weakly advantageous mutant with *s* = 10^−2^ by varying the initial condition *ω*(*x*). For weakly advantageous mutants, the theory is well approximated by *ρ*(*s, x*) ≃ *sN*_i_*Z*_t_*ω*(*x*). Blue circles denote simulation results; black lines represent the corresponding fixation probability on a fully connected graph of size *N*_i_; the gray line indicates the fixation probability of a neutral mutant (1*/N*_i_); and the blue line shows theoretical prediction. Other parameters are *β* = 6 × 10^−4^ and *γ* = 3 × 10^−4^. For each parameter set, fixation probabilities were estimated from 2 × 10^5^ independent simulations.

The result for the advantageous mutants is consistent with that of [6]; for disadvantageous mutants we find an even stronger effect: the fixation probability of a mutant is increased by orders of magnitude relative to the fully connected case.

Here we develop an analytical approximation for the fixation probability. The main difficulties are that fixation occurs within a heterogeneous network [12] and inside an infected subpopulation that fluctuates over time and space. To reduce complexity in the analysis while retaining degree heterogeneity, we approximate the fluctuating infection state of each node *x*, that is, whether the node is infected or susceptible, by the probability *p*_*x*_ of being infected at steady state. Thus, the temporally fluctuating infected subpopulation is replaced by a static infection-probability profile defined over all nodes of the network (Appendices A 3 & B 1).

Within this approximation, the dynamics become simpler because we no longer track the time-dependent infection state of each node. Rather, we ask a different question: Given that node *x* is infected with probability *p*_*x*_, what type of infection does it carry? This infection type is tracked by the indicator *η*(*x*), which is one if node *x* carries a mutant and zero otherwise. We derive the transition probabilities for changes in *η*(*x*), and then identify a martingale of the process whose ensemble average remains invariant under the dynamics:

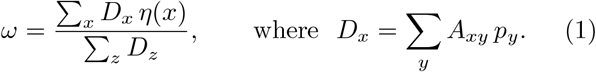

The quantity *D*_*x*_ measures the average number of wild-type infected neighbors of node *x* at the wild-type recovery steady state. The martingale *ω* is the normalized sum of *D*_*x*_ over nodes carrying a mutant. Thus, if a single mutant is initially located at node *x*, only that node contributes, and *ω* becomes a function of the initial mutant location, *ω*(*x*) = *D*_*x*_/Σ_*z*_ *D*_*z*_ (Appendices B 1 & B 2).

In the weak-selection limit, the ratio of the drift *E*(Δ*ω*) to the diffusion *E*(Δ*ω*^2^*/*2) of *ω*(*x*) determines the initial-condition-dependent fixation probability *ρ*(*x, s*) of a mutant initially introduced at node *x*. From this, we obtain the fixation probability *ρ*(*s*) of a mutant that arises randomly on a wild-type infected node. This theory works very well for the star-graph systems, and it reproduces the same order of suppression of selection, as shown by the blue lines in FIG. 1(a-d) (Appendices B 3 and B 4).

A qualitative mechanism of “hub-holding” was proposed to explain selection suppression in a star graph [6]. In a star graph, the hub dominates transmission and remains infected most of the time, so most transmission events simply copy its state to leaves and are effectively neutral. Selection acts only during rare hub re-infection events, when competing leaf types determine which type occupies the hub. Thus, the predominance of neutral transitions suppresses selection; we posit that the same mechanism operates in scale-free networks.

Here, we derive a quantitative measure of selection suppression.

The observations (FIG. 1(a-d) & [2, 6]) motivate a *model-derived* measure of hub dominance that quantifies the factor by which selection is suppressed. The *suppression factor*,

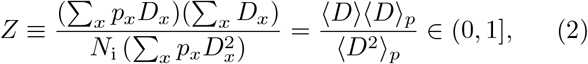

appears in the expression for *ρ*(*s*) (Eq. B29) as a multiplicative factor modifying the strength of selection, with smaller *Z* indicating stronger suppression. Here 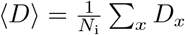 is the expected infected-neighbor count of a node, 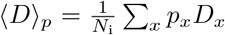 is the expected infectedneighbor count of an infected node, 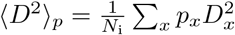 and *Z* is an inverse measure of heterogeneity across the network, which decreases in response to the presence of hubs.

We estimate *Z* in two ways: Analytically, by deriving the stationary probabilities *p*_*x*_ (Appendix A 3) and substituting them into Eq. (2), and numerically, from simulations (Appendix B 4). We denote these estimates by *Z*_t_ and *Z*_s_, respectively. Although the theory gives a slight systematic underestimation of selection suppression as selection strength increases (FIG. 1(a-d)), in the weak-selection limit it captures both the average fixation probability *ρ*(*s*) and the initial-condition-dependent fixation probability *ρ*(*s, x*) (FIG. 1(e)). This agreement also makes explicit the role of *Z* as the factor suppressing the strength of selection. In the case of disadvantageous mutants, selection suppression leads to a higher selection-mutation balance of the mutant infection on a heterogeneous networks compared to that on a degree-homogeneous network with the same infected population size, *N*_i_, see FIG. 8(b).

The suppression factor in Eq. (2) depends on the parameters of infection through the quantity {*p*_*x*_}. In the limit *β/γ* → ∞, the ratio reduces to *Z*_A_ = ⟨*k*⟩ ^2^*/* ⟨*k*^2^⟩, a structural property of the network (Appendix B 3). Note that *Z*_*A*_ ≤ *Z*, that is, the stronger the infection, the stronger the suppression effect.

Therefore, *Z*_*A*_ measures the maximal strength of suppression that characterizes a given network. This raises a natural question: which networks minimize *Z*_A_ and therefore most strongly suppress selection? Within the family of large scale-free networks, selection suppression is strongest near a power-law exponent of two of the degree-distribution: both heavier and thinner tails increase *Z*_A_ and therefore weaken the suppression. Another interesting case is a nearly regular network (of degree *k*_1_) with a small fraction of highly connected hubs with a higher degree, *k*_2_. For this type of networks, maximal suppression occurs at the degree-balanced two-point distribution, where the probability of each degree class is inversely proportional to its degree, so that each class contributes equally to the mean degree: *p*_1_*k*_1_ = *p*_2_*k*_2_, where *p*_*i*_ is the probability to have degree *k*_*i*_ (Appendix B 5). In general, strong suppression requires a degree distribution with nonzero probability at both low and high degrees, such that, while low-degree nodes are abundant, high-degree hubs remain sufficiently influential.

We next use *Z*_A_ to estimate degree-heterogeneity in empirical networks of social behaviors important for disease spread, including direct physical contact and proximity. Analyzing more than a thousand animal social networks from the animal social network repository [13], we find that most networks have *Z*_A_ *<* 1, with many characterized by a low structural selection suppression factor, *Z*_A_ (FIG. 2). Thus, empirical social networks are sufficiently heterogeneous for selection suppression to be a ubiquitous feature of real world communities, rather than a peculiarity of synthetic heterogeneous networks (see also FIG. 6).

**FIG. 2.**
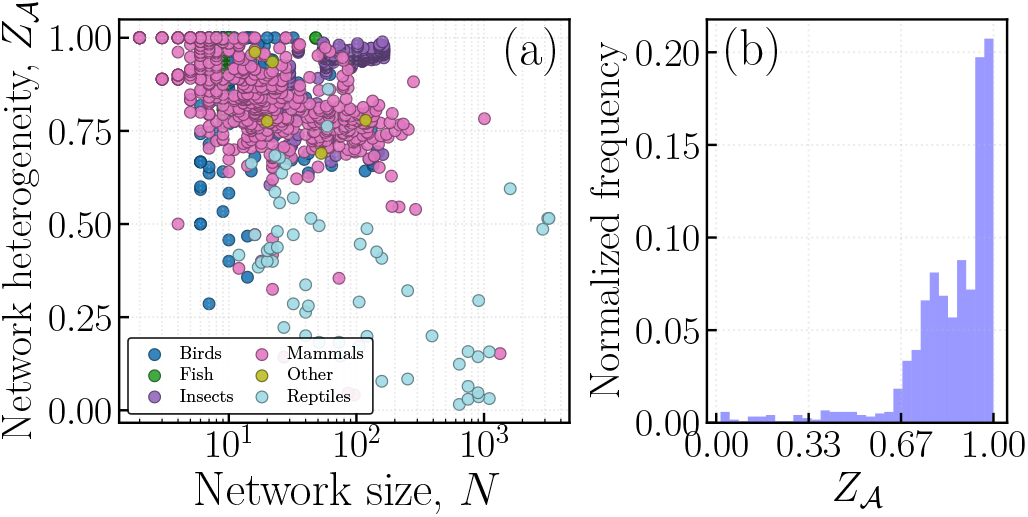
Degree heterogeneity and suppression of selection in empirical networks. (a) All networks from the repository [13] are projected onto the *Z*_*A*_–*N* plane. (b) The histogram shows the normalized frequency of networks from the repository within each bin of *Z*_*A*_. We used thirty equal-width bins across the full range of *Z*_*A*_.

**FIG. 3.**
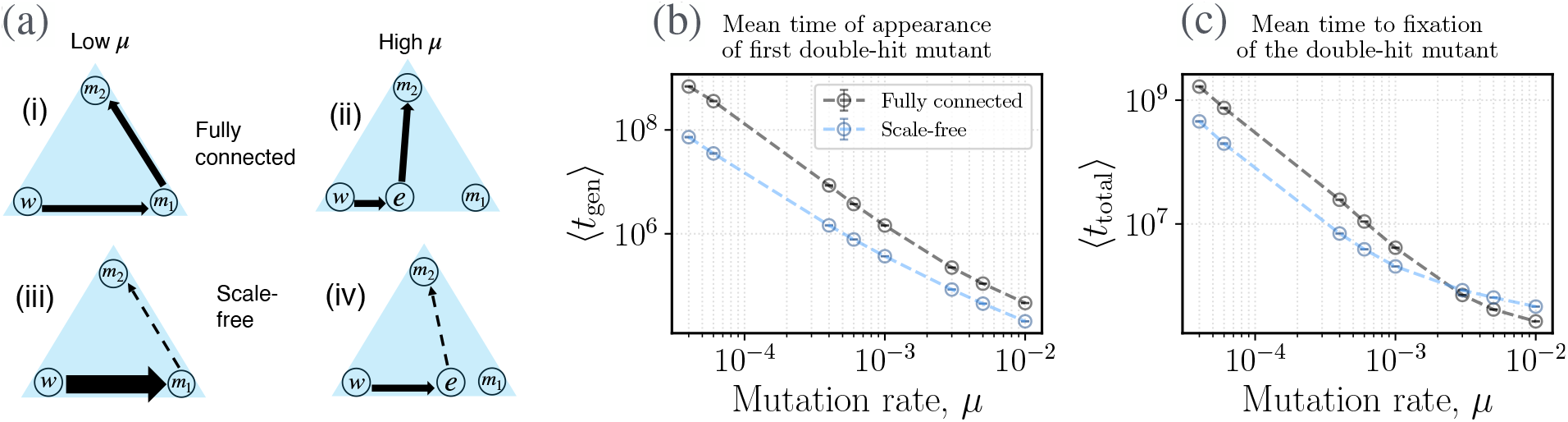
Evolutionary consequences of suppression of selection on degree-heterogeneous networks. Panel (a) qualitatively compares fitness-valley crossing in the low- and high-mutation regimes on fully connected and scale-free networks. The long-lived states are shown on a simplex: *w* is the all-wild-type state, *e* is the mutation–selection balance between wild type and single-hit mutants, *m*_1_ is the all-single-hit-mutant state, and *m*_2_ is the all-double-hit-mutant state. Arrows indicate transitions between states, with thick solid, thin solid, and dashed arrows denoting high, intermediate, and low transition rates, respectively. Panels (b) and (c) compare the mean generation time *t*_gen_ and the total time to fixation, including the generation of the double-hit mutant, for fully connected (gray) and scale-free (blue) networks, respectively. In this case, the fully connected and scale-free (Barabasi–Albert) networks have (*N, β*) = (1.456 × 10^3^, 3.7 × 10^−7^) and (10^3^, 6 × 10^−4^), respectively, with the same recovery rate *γ* = 3 × 10^−4^. This parameter choice gives the same infected subpopulation size, *N*_*i*_ ≈ 651, and basic reproductive number, *R*_0_ = 1.81, for both networks. The single-hit and double-hit mutants have infection rates that are 10% lower and 50% higher than those of the wild-type, respectively. The mutation rates from wild-type to single-hit mutants and from single-hit to double-hit mutants are identical, *µ* = 4 × 10^−5^. Each data point in (b,c) is obtained from at least 3.2 × 10^3^ independent simulations. Standard error bars are extremely small and are barely visible.

Apart from mutant fixation probability and selection-mutation balance, fixation suppression affects the rate of evolution of complex phenotypes. Here we consider *fitness-valley crossing* [14–19], where the observed asymmetric suppression of advantageous and deleterious selection can have nontrivial consequences. Fitness-valley crossing occurs when the single-hit mutation is deleterious, but a *double-hit* mutation arising from the single-hit mutant creates a strongly advantageous genotype which takes over the population. We compare the generation time *t*_gen_ and total fixation time *t*_total_ of the double-hit mutant on fully connected and scale-free networks with identical *R*_0_ and *N*_i_. For simplicity, the wild-type to single-hit mutation rate and the single-hit to double-hit mutation rate are both set to *µ*.

Let *w, m*_1_, and *m*_2_ denote the three pure states of the system, consisting entirely of wild type, single-hit mutants, and double-hit mutants, respectively. The mutation–selection balance between wild type and singlehit mutants is represented by *e*. Two routes to fitness-valley crossing are possible: Sequential fixation, P_seq_ : *w* → *m*_1_ → *m*_2_, in which the single-hit mutant fixes before the double-hit mutant, and stochastic tunneling, *P*_tun_ : *w* → *e* → *m*_2_, in which the double-hit mutant arises from the mutation–selection balance *e* and fixes without the system reaching *m*_1_.

On a fully connected network with very low *µ*, the dominant path is *P*_seq_, and for high-*µ* cases, *P*_tun_ dominates [14, 20]. However, asymmetric modulation of disadvantageous and advantageous selection on heterogeneous networks changes the rates of these paths. For the low-*µ* cases where the path_tun_ is nearly impossible, single-hit mutant fixation gets enormous boost on heterogeneous networks. A quickly fixated single-hit mutant can lower *t*_gen_ and *t*_total_, thereby accelerating fitness-valley crossing. For high-*µ* cases, where *P*_tun_ dominates, the enhanced fixation probability of the single-hit mutant becomes immaterial; although heterogeneity increases the mutation–selection balance and can reduce *t*_gen_, the suppressed selection of the advantageous double-hit mutant increases *t*_total_ relative to the fully connected network, slowing fitness-valley crossing. We conclude that, when *N*_i_*µ* ≪ 1, degree heterogeneity can significantly accelerate the adaptation of complex traits.

Previous work demonstrated that degree heterogeneity, and the presence of super-spreading individuals in particular, can have a major influence on the basic reproductive number of a pathogen and can substantially enhance the rate of infection spread [3]. Numerical simulations of the evolution of advantageous pathogen mutants subsequently showed that superspreaders can at the same time slow down the emergence and invasion of advantageous pathogen mutants, brought about by a hub-holding effect, which weakens selection [6]. Here, we build on this work and developed mathematical theory that describes the effect of degree heterogeneity in networks on pathogen evolution. We find that selection is weakened even more strongly for deleterious mutants, which behave almost like neutral mutants in the presence of strong degree heterogeneity. Based on our theory, we defined network characteristics that lead to this effect, and found that a variety of empirically derived animal interaction and association networks display these characteristics. These findings have important implications for understanding pathogen evolution. On a basic level, our results mean that the presence of superspreaders in social networks can lead to a large burden of deleterious pathogen mutants in a population, orders of magnitude larger than expected by established evolutionary theory. This has further implications for the rate of adaptive evolution. Complex traits require the accumulation of more than one mutation in the same individual, and frequently, intermediate mutants carry a fitness cost, thus requiring the crossing of a fitness valley [14–19]. We have shown that under relatively low mutation rates, the presence of superspreaders can substantially increase the rate at which a multi-hit mutant is produced, and can further shorten the time until the multi-hit mutant invades the population. Although superspreaders slow down the spread of advantageous mutants by themselves, as previously reported [6], the reduced selection against disadvantageous mutants is the stronger effect, thus accounting for the observation that superspreaders can accelerate adaptive evolution. In conclusion, we present an analytical understanding of pathogen evolution on networks with degree heterogeneity, and present new results showing the importance of superspreaders for the maintenance of deleterious mutants, and for accelerating adaptive evolutionary processes that build on this reservoir of deleterious mutants.

In the current paper, we focused exclusively on super-spreading individuals that are characterized by a relatively high degree of social connectivity. In addition to this, pathogen superspreading can be brought about by other mechanisms, such as the presence of selected host individuals in a population who are more prone to infection and to pathogen transmission (e.g. due to genetic heterogeneity). The types of social behaviors underlying network structure (e.g., direct contact vs. spatial proximity) also drive transmission dynamics, and could further impact the role of superspreaders and pathogen evolution. It will be interesting to investigate how such different modes of super-spreading affect pathogen evolution, and how this compares to the dynamics reported here.

This work was supported by NSF grant DMS 2424853 (DW and NK) and NSF grant DMS 2435484 (NK and DW). SSM gratefully acknowledges fruitful discussions with Ramanarayanan Kizhuttil and Sydney Ackermann.

## Appendix A Model formulation & simulation

### 1. Infection process

Consider a population of *N* individuals, where each individual can exist in one of two states, infected or susceptible. Each infected individual can interact and transmit infection only to a limited subset of the population rather than to all others. Such interactions are naturally represented by a *network*, in which nodes correspond to individuals and edges denote possible transmission routes. Infection can spread only from an infected node to its immediate neighbors, and the probability of transmission depends on how frequently these neighbors interact. These interaction frequencies are encoded in the *adjacency matrix A* = (*A*_*xy*_), where each entry *A*_*xy*_ ∈ [0, 1] represents the relative frequency of contact between individuals *x* and *y*. A value of *A*_*xy*_ = 1 indicates a pair that interacts with maximal frequency, whereas *A*_*xy*_ = 0 implies no direct interaction between them. Here, we focus on undirected networks, i.e., *A*_*xy*_ = *A*_*yx*_ for all pairs. Thus, if node *x* can transmit infection to node *y*, the reverse is also possible. The *degree* of node *x*, defined as *k*_*x*_ = ∑_*y*_ *A*_*xy*_, measures how frequently node *x* interacts with others and therefore indicates its potential to spread infection. The infection dynamics is driven by two opposing processes—*transmission* and *recovery*. An infected node *x* can transmit infection to its susceptible neighbors at a rate *β* per interaction. Since the likelihood of interaction scales with *k*_*x*_, the total rate at which node *x* attempts to transmit infection is *βk*_*x*_. However, recovery occurs independently of the network structure: Each infected node returns to the susceptible state at a constant rate *γ*.

### 2. Simulation

We simulate infection process on a weighted scale-free network of size *N*. To characterize the baseline infected subpopulation, we first evolve a population consisting entirely of wild-type infections. Each node *x* is in one of two states: Susceptible or infected. At every update step, we identify all infected nodes, and each infected node performs one stochastic event on average. For an infected node *x*, its weighted degree is *k*_*x*_ = ∑_*z*_ *A*_*xz*_, where *A*_*xz*_ denotes the weight of the edge between nodes *x* and *z*. With probability *β*_w_*k*_*x*_, node *x* attempts to infect one of its neighbors, chosen with a probability proportional to the corresponding edge weights. If the selected neighbor is susceptible, it becomes infected; otherwise the attempt is aborted. With probability *γ*, node *x* recovers and becomes susceptible.

The system is evolved for a burn-in period until it reaches an infection–recovery quasi-steady state. Then, a single mutant is introduced by converting one randomly selected infected node into the mutant type, which spreads with infection rate *β*_m_. From this point onward, the stochastic dynamics lead to one of two possible outcomes: the mutant strain fixates within the infected sub-population, or it goes extinct, leaving the wild-type infected subpopulation unchanged. By repeating this process over a large number of independent realizations, we estimate the mutant fixation probability *ρ*(*s*) from simulation.

For the mutation–selection balance and valley-crossing simulations, we consider a constant mutation rate *µ*. With probability *µ*, a newly infected individual mutates at the time of infection and acquires a mutated infection type instead of inheriting the infection type of its source.

### 3. Steady-state infected subpopulation

Throughout this work, we assume that the ratio *β/γ* lies beyond the *epidemic threshold* determined by the network *A*, ensuring a persistent nontrivial endemic steady state. Let *p*_*x*_ ∈ [0, 1] denote the probability that node *x* is infected. The dynamics of the infection probability *p*_*x*_ are governed by two competing processes. First, an uninfected node *x* can become infected by contacting its infected neighbors. The gain rate of infection at *x* is

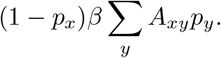

Second, an infected node *x* can recover at a constant rate *γ*, giving the loss term

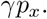

At steady state, the gain and loss terms balance, yielding the condition

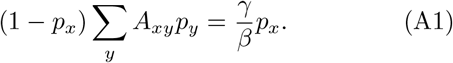

In the deterministic limit, where stochastic fluctuations in the infection fraction are negligible, the time averaged *size of the infected subpopulation* is given by

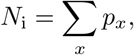

which quantifies the expected number of infected individuals at steady state. Because recovery continually returns nodes to the susceptible state, this effective size remains smaller than the total population size *N*. For a fully connected network, the steady-state infection probability takes the form

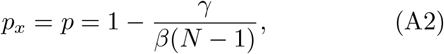

so that

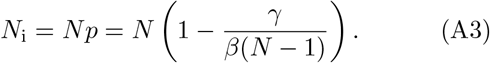

For a fully connected graph, the infectivity and the size of the steady-state infected subpopulation are related by

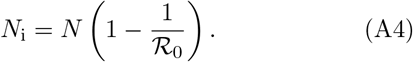

Where, *R*_0_ is the basic reproductive number. Comparing Eqs. (A3) and (A4), we obtain the following expression for the basic reproductive number

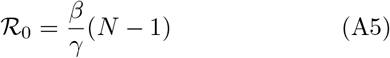

for a fully conncted network.

### 4. Fixation of a mutant pathogen in an infected subpopulation at a quasi-steady state

Consider a population in which an infection with transmission rate *β*_w_ = *β* has already established a quasisteady endemic state, maintained by the balance between transmission and recovery. Into this steady-state background, a new infection is introduced at a single node. The new strain differs slightly in its transmission ability, characterized by a modified rate

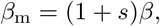

where the dimensionless parameter *s* quantifies the relative advantage or disadvantage of the mutant infection in transmitting to susceptible individuals. A positive *s* corresponds to a more transmissible strain, while a negative *s* indicates a less transmissible one. Throughout this analysis, we focus on the weak-selection case, |*s*| → 0, where the perturbation introduced by the new infection is small compared with the steady-state dynamics of the resident strain. In this limit, the overall size of the infected subpopulation and the nodal infection probabilities *p*_*x*_ remain approximately constant, allowing the dynamics of the new infection to be studied on a fixed background of endemic infection. Our objective is to determine how the structure of the underlying contact network modulates the likelihood that the newly introduced infection either goes extinct or successfully replaces the resident strain, and how these fixation probabilities scale with the relative transmission advantage *s*.

## Appendix B Approximated process

### 1. A simplified model and the martingale

Here we propose a simplified process which makes it possible to identify a conserved quantity essential for this study. We then demonstrate that the results are relevant for the original process considered in the main text.

To simplify the description of infection dynamics, we assume that each node is effectively always infected, but its contribution to transmission is weighted by its steady-state infection probability *p*_*x*_. The stochastic infection process is then described by sequential update network states *η* = {*η*(*x*)}, where *η*(*x*) ∈ {0, 1} denotes the infection type at node *x*. Here, *η*(*x*) = 0 represents the wild-type infection, and *η*(*x*) = 1 represents the mutant infection. At each update step, a node *x* is selected for state replacement with probability proportional to its steady-state infection probability *p*_*x*_, that is,

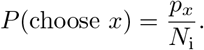

One of its neighbors *y* is then selected as the source of infection. We assume that neighbor *y* transmits infection at a rate proportional to its degree and its intrinsic infection rate. Thus, the probability that *y* is chosen as the infector and transmits to its randomly chosen neighbor *x* is given by *A*_*xy*_*p*_*y*_*/* ∑_*z*_ *A*_*xz*_*p*_*z*_. Consequently, the total transition probability that node *x* changes its infection state, given the configuration *η*, is then written as

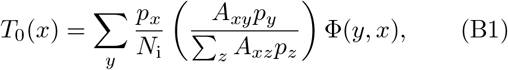

where Φ(*x, y*) = *η*(*x*) [1 − *η*(*y*)] + *η*(*y*) [1 − *η*(*x*)], and *η*^*x*^ denotes the network configuration after the state change has occurred at node *x*. Note that the information 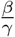 is incorporated in the quantities {*p*_*x*_}. The average mutant density is defined as

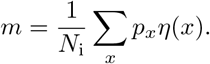

When node *x* flips its state the resulting change in *m* is given by

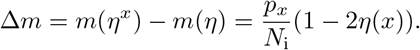

Expected change in *m* after one update step is obtained by averaging over all nodes, weighted by their transition probabilities as follows

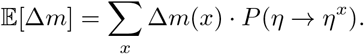

Which is a non-zero quantity in general. We seek a quantity that depends on the node state *η*(*x*) and remains conserved. We propose the following quantity [21]

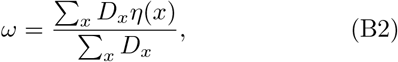

is conserved. Here *D*_*x*_ = ∑_*z*_ *A*_*xz*_*p*_*z*_. If node *x* changes its state, the resulting change in this quantity is

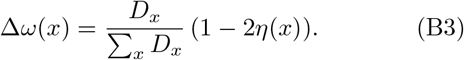

To check its conservation, we compute the expected change in this quantity after a flip at node *x*. The expected change is written as

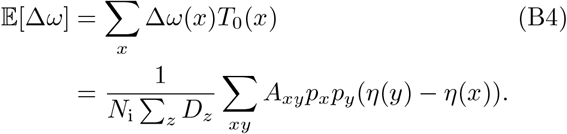

By symmetry of *A*_*xy*_ and exchange of dummy indices, the numerator vanishes, hence

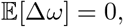

confirming that *ω* is conserved when mutant is neutral.

### 2. Test of conservation of *ω*

To validate the conservation of the quantity *ω* under neutral dynamics, we performed simulations of the infection process on a scale-free network and explicitly tracked the temporal evolution of *ω* after mutant introduction. We first generated a scale-free network using a preferential-attachment algorithm. The system was initialized with a population consisting entirely of wild-type infections and evolved under infection–recovery dynamics until a steady state was reached. From this steady state, we computed the nodal infection probabilities *p*_*x*_ by time-averaging the infection status of each node. These probabilities define the effective weights entering the conserved quantity *ω*.

After the burn-in period, a predetermined subset of nodes chosen among those with the highest infection probabilities was designated as potential mutant introduction sites. Mutants were introduced only at nodes within this subset that were infected at the time of introduction, ensuring that mutants arise exclusively within the infected subpopulation, in epidemiologically relevant locations, and without significantly perturbing the system. The system was then evolved forward in time under neutral dynamics. At each time step following mutant introduction, we recorded both the weighted mutant density *m* and the conserved quantity *ω*.

The variable *η*(*x*) appearing in the definitions of *ω* and *m* requires careful specification. While the full infection-process simulation involves three possible states for each node—wild type, mutant, and susceptible—our approximated description treats *η*(*x*) as a binary variable that records infection lineage rather than instantaneous infection status. If node *x* is currently infected, *η*(*x*) is determined directly by its present infection state: *η*(*x*) = 1 for a mutant infection and *η*(*x*) = 0 for a wild-type infection. If node *x* is in the recovered (susceptible) state, *η*(*x*) is assigned according to the identity of the most recent infection carried by that node prior to recovery.

We repeated this procedure over many independent realizations and averaged the resulting trajectories. While the averaged mutant density remains time dependent, the quantity *ω* stays nearly constant in time, confirming its approximate conservation under neutral dynamics (see FIG. 4). This numerical test provides direct support for the analytical conservation law derived in Appendix B 1.

**FIG. 4.**
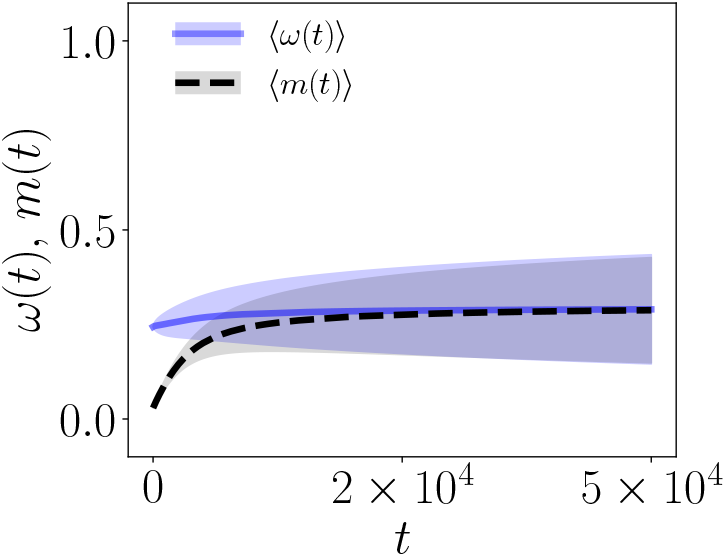
Numerical verification of conservation of ensemble averaged ω. Time evolution of the ensemble averaged conserved quantity *ω* (blue) and the weighted mutant density (black dashed) following neutral mutant introduction on a scale-free network. While the mutant density *m* changes due to the drift, *ω* remains nearly constant over time (initially drifting slightly, but rapidly converging to a constant value), confirming its approximate conservation under neutral dynamics. Total 3.2 × 10^3^ independent trajectories are used to get the average; Transparent envelop represents standard deviation.

### 3. Evolution of the conserved quantity under weak selection (*s* → 0)

In the presence of selection, the transition probability (that is, the probability that node *x* changes its infection state) takes the following form

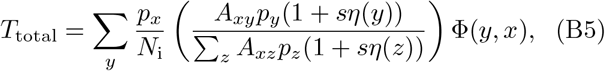

where *s* denotes the selection strength. In the limit of weak selection, we can approximate the fitness ratio as

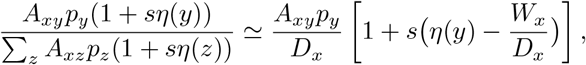

where

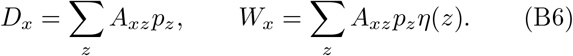

After rewriting the transition probability using the weak selection approximation, we can separate it into two parts, a neutral part *T*_0_ and a selection part *T*_*s*_:

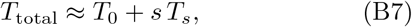

where

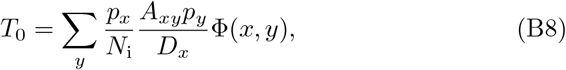

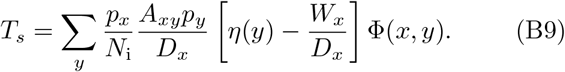

The first term *T*_0_ corresponds to the neutral dynamics, while *T*_*s*_ represents the correction due to weak selection. We can write the expected change in *ω*(*x*) as follows

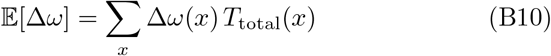

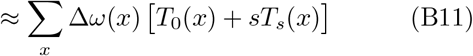

As we know, the expected change in *ω* receives a nonzero contribution only from the *T*_*s*_ term (we proved that the *T*_0_ terms do not contribute, see Eq. (B4)). So the expected change in *ω* takes the form as follows

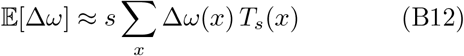

Now let us substitute Δ*ω* and *T*_*s*_ from Eqs. (B3) and (B9) into Eq. (B12), and we obtain the following equation,

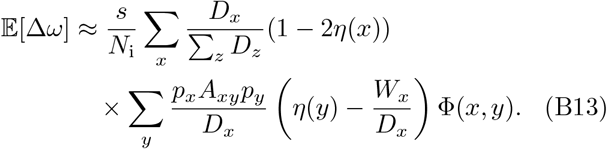

Using the identity (1 − 2*η*(*x*))Φ(*x, y*) = (*η*(*y*) − *η*(*x*)) in above equation we get

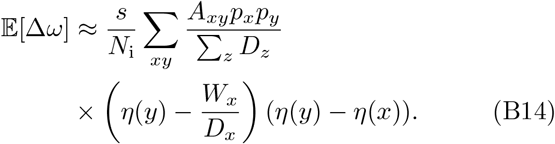

In Eq. (B14), the term *W*_*x*_*/D*_*x*_ represents the fraction of infected neighbors of node *x* that are mutants, i.e., a local mutant density around node *x* among the infected neighbors. Under the mean-field assumption for degree uncorrelated large network [21], fluctuations of *η*(*x*) and *η*(*y*) are sufficiently fast and weakly correlated across nodes. As a result, the local mutant density 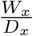 becomes effectively uncorrelated with the difference term (*η*(*y*) − *η*(*x*)). Therefore, their expectation factorizes as

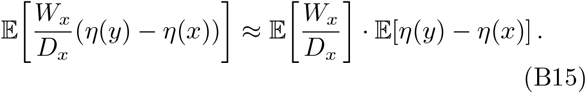

By symmetry of the mean-field dynamics, we have

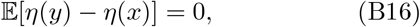

which implies

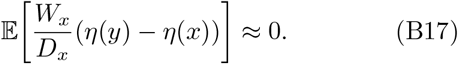

Hence, the contribution of this term in Eq. (B14) vanishes in the mean-field limit. Hence we can rewrite Eq. (B14) as follows

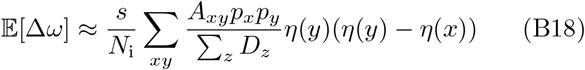

Now, let us focus on the term *η*(*y*) *η*(*y*) − *η*(*x*) appearing in the previous equation. This term makes the summation intractable. However, we can simplify it using a mean-field approximation as follows

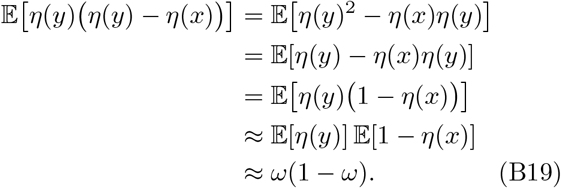

Using approximation (B19) in Eq. (B14) we get

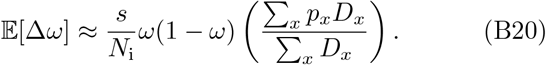

Let us now compute the expected value of (Δ*ω*)^2^. We note that the evaluation of this term does not require selection, since the diffusion arises from stochastic fluctuations alone. Therefore, the diffusion term can be approximated using the neutral-case transition probability *T*_0_. Squaring Eq. (B3), we obtain

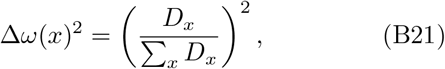

expected value of which can be computed as

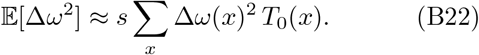

Substituting Δ*ω*(*x*)^2^ and *T*_0_ from Eqs. (B3) and (B8) into Eq. (B22), we obtain

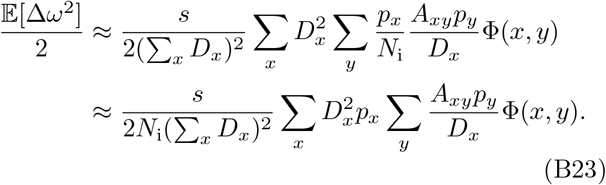

Now, let us focus on the term Φ(*x, y*) = *η*(*y*)(1−*η*(*x*)) + *η*(*x*)(1 − *η*(*y*) appearing in the previous equation. This term makes the summation intractable. However, we can simplify it using a mean-field approximation as follows

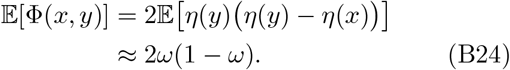

Using approximation (B24) in Eq. (B23) we get

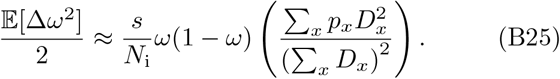

The ratio of the drift to the diffusion term, *α*, is given by

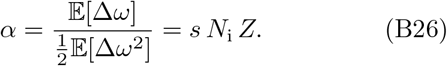

Here *Z* denotes the *suppression ratio*,

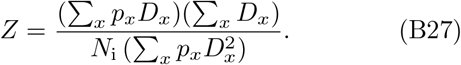

In the high-infectivity limit *γ/β* → ∞, where *p*_*x*_ → 1 for all *x, N*_i_ → *N*, and *D*_*x*_ = *k*_*x*_, this expression reduces to

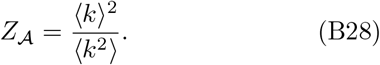

We note that *Z*_A_ is determined solely by the underlying network, while *Z* is a function of both the network structure and the infectivity. Having obtained the ratio *α*, one can, in principle, directly express the position-dependent fixation probability using Eq. (B29), as shown in [21].

### 4. Fixation probability

Having determined the conserved quantity along with its drift and diffusion coefficients, we can express the fixation probability of a mutant introduced at an infected node *x* as [21],

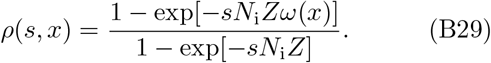

Since each node has a different probability of being infected by the wild type in the first place, the appearance of mutants should be conditioned on that probability. Therefore, the average fixation probability of a mutant can be written as

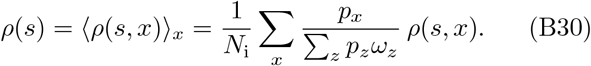

In the limit *s* → 0, we have

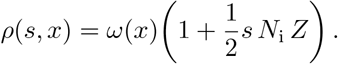

Therefore,

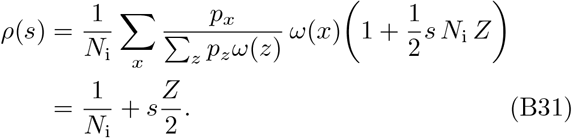

The slope of the fixation–probability curve at *s* = 0 is *Z/*2. We can compute *Z* in two different ways: *Z*_t_ is obtained analytically from our theoretical expression in Eq. (2), while *Z*_s_ is computed directly from simulation data by estimating the slope of the fixation–probability curve *ρ*(*s*)_s_ in a small neighborhood around *s* = 0. The slope of the fixation–probability curve at *s* = 0 is *Z/*2, and therefore

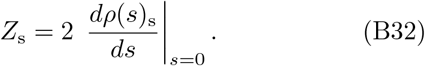

In the main text, we compare the fixation probability of a mutant arising in an infection process with the fixation probability of a mutant in a Moran process on a fully connected graph. The fixation probability in the fully connected case is given by [1]

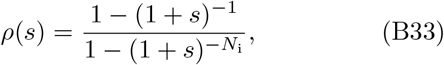

where *s* denotes the selection strength and *N*_i_ is the size of the infected subpopulation. We note that, for an infection process on any homogeneous-degree (regular) graph with infected subpopulation size *N*_i_, the mutant fixation probability is the same as that of the Moran process on a fully connected network of size *N*_i_, as given by the above equation. This is because, in regular graphs, degree heterogeneity is absent and therefore does not generate amplification or suppression of selection.

### 5. Maximization of suppression of selection

*Two point degree distribution:* To gain analytical insight into the effect of degree heterogeneity on

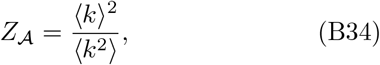

we consider a simple two–point degree distribution. In this case, nodes can have only two possible degrees, *k*_1_ and *k*_2_, with probabilities *p* and 1 − *p*, respectively, so that

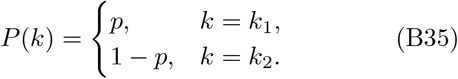

Let *r* = *k*_2_*/k*_1_ denote the ratio between the two degrees. The first and second moments of the degree distribution are ⟨*k*⟩ = *pk*_1_ + (1 − *p*)*k*_2_ and 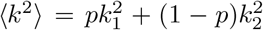, which lead to

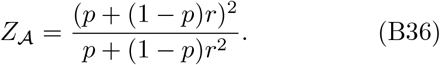

Minimizing *Z*_A_ with respect to *p* for fixed *r* yields the optimal probability

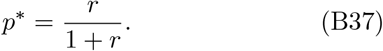

Consequently, the remaining fraction of nodes with degree *k*_2_ is

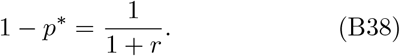

Thus, there is an optimal weight on the tail of the distribution, below and above which the *Z* decreases.

#### Scale-free degree distribution

We next consider a scale–free degree distribution of the form

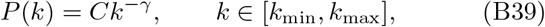

where *γ >* 1 is the degree exponent and *C* is a normalization constant. Normalizing the distribution gives

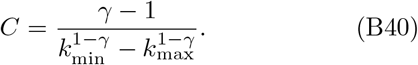

The moments of the distribution are

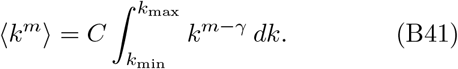

Evaluating these moments shows that *Z*(*γ*) attains a minimum near *γ* ≈ 2. Thus, strong suppression arises for heavy-tailed degree distributions, but diminishes when the tail becomes too extreme.

### 6. Estimation of R_0_ for a finite scale-free graph

For a large single-component graph, a closed-form expression for the basic reproductive number already exists in the literature [8] and can be written as

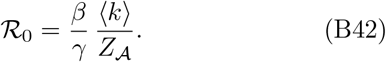

For large Albert–Barabási networks, this quantity diverges [8]. We consider a finite Barabási–Albert graph and compute *Z*_A_ numerically. We start with an infection-free host network, select a source node, introduce a single infection at that node, and compute the average number of secondary infections generated before the source node recovers. We then repeat this procedure for all nodes in the network to obtain the network-averaged basic reproductive number R_0_.

## Appendix C Effect of increasing infectivity on a star graph

Given the infection rate *β* and recovery rate *γ*, we can obtain a theoretical estimate of the suppression factor *Z*. For a star graph of size *N*, the central node remains infected for the majority of the time. Therefore, we approximate the infection frequency of the center as *p*_c_ ≈ 1. The infection frequency of a leaf node can be approximated as

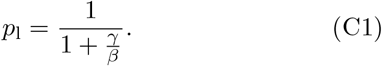

With all other parameters held fixed, increasing *β* results in a larger infected subpopulation size by increasing the infection frequencies of leaf nodes, *p*_l_. Using these frequencies, the suppression factor *Z* can be estimated from Eq. (2) as

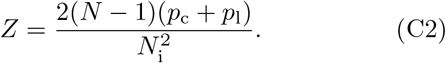

In the limit of large *β*, the infected subpopulation size *N*_i_ approaches *N*, and the suppression factor simplifies to

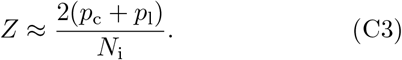

Thus, the suppression factor roughly scales as *Z* ∼ 1*/N*_i_. Substituting this into Eq. (B31), the product of the fixation probability and the infected subpopulation size becomes

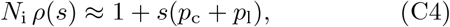

which is approximately independent of *N*_i_. In contrast, for a star graph of size *N*_i_ under the standard Birth–death process, the fixation probability satisfies

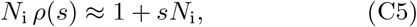

which is clearly an increasing function of *N*_i_. FIG. 5 illustrates this. Keeping the recovery rate fixed, increasing the infection rate naturally leads to an increase in the infected subpopulation size. In a star graph, a higher infection rate increases the number of infected leaf nodes, thereby resulting in a larger effective infected subpopulation size *N*_i_. In the weak-selection limit, the fixation probability can be written as 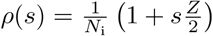, which implies that the product 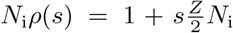. When this product is plotted as a function of *N*_i_, obtained by varying *β*, it remains approximately constant. This observation indicates that the suppression factor scales as *Z* ∼ 1*/N*_i_ on a star graph. Therefore, increasing infectivity leads to a larger infected subpopulation size *N*_i_, a corresponding decrease in the neutral fixation probability 1*/N*_i_, and a more pronounced suppression effect through a reduced value of *Z*. Whereas, for a mutant in the birth–death process on a star graph of size *N*_i_, this product varies as 1 + *sN*_i_, reflecting the fact that the suppression ratio is *Z* = 2, corresponding to amplification of selection and independence from the infected subpopulation size.

**FIG. 5.**
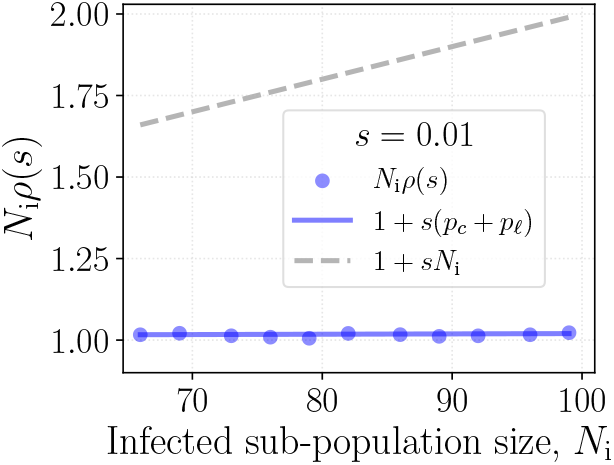
Effect of increasing infectivity on a star graph of fixed size. The total population size is fixed at *N* = 10^2^. Other parameters are set to *s* = 0.01 and *γ* = 3 × 10^−4^. The infectivity parameter *β* is varied to generate different effective infected subpopulation sizes *N*_i_, as shown in the plot. We display the product of the infected population size and the fixation probability as a function of the infected subpopulation size. The gray curve represents the case of a mutant under the Birth– death process for a star graph of the corresponding size. The blue scatter points are obtained from simulations of the infection process, while the blue solid line denotes the theoretical prediction for a star graph that incorporates the suppression factor *Z*.

**FIG. 6.**
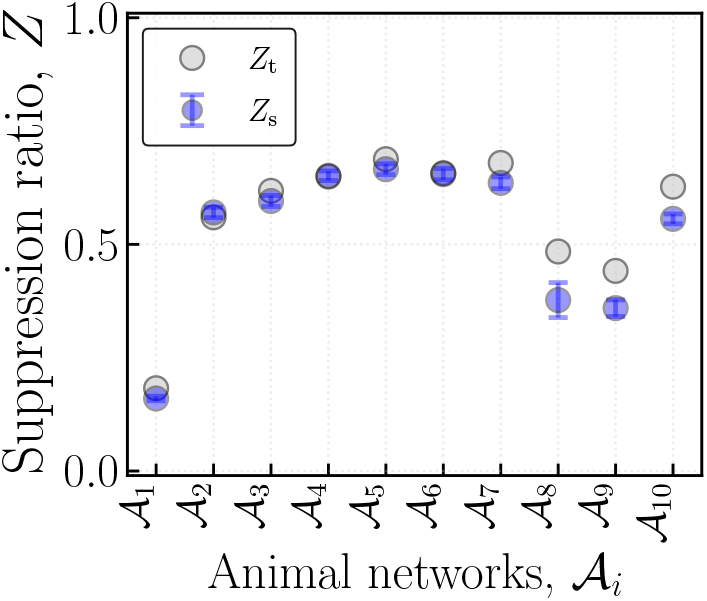
Degree heterogeneity and suppression of selection in empirical networks. Comparison between the theoretical suppression factor *Z*_t_ and the simulation-based suppression factor *Z*_s_ for the ten selected networks listed in TABLE I. Simulation values include propagated standard deviations shown as error bars. Parameters: *β* = 6 × 10^−4^ and *γ* = 3 × 10^−4^.

**FIG. 7.**
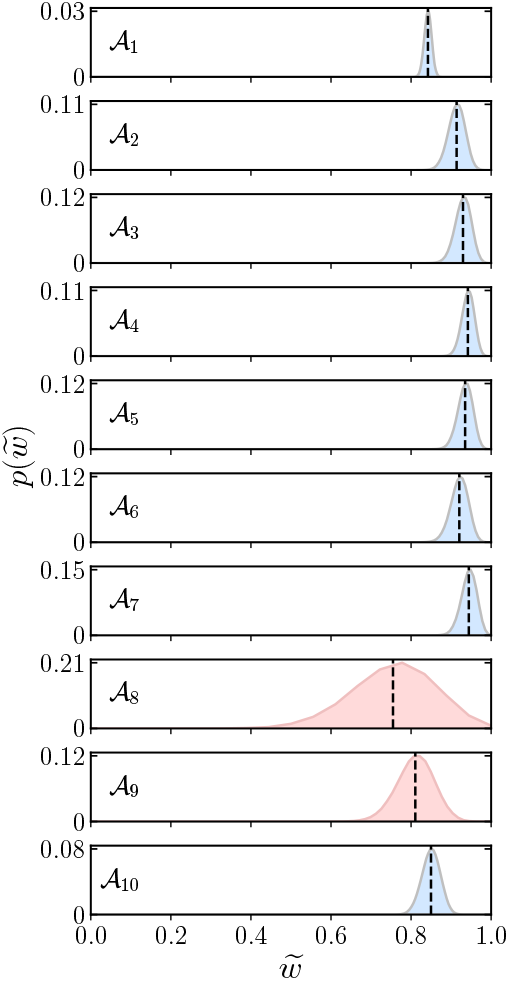
Fluctuations in the density of infection at the wildtype-recovery steady state across all networks listed in TABLE I. We analyze ten empirical networks (*A*_1_– *A*_10_). For each network, once the system reaches the wildtype-recovery equilibrium, we recorded the total number of infected nodes normalized by the network size, 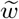, over time. For most networks, the probability density 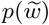 remains narrow, with the majority of the probability mass concentrated around the mean represented by dashed vertical lines (blue curves). However, a subset of networks exhibits significantly broader distributions (red curves), indicating enhanced fluctuations around the average infection level. These networks can therefore be identified as outliers with substantially larger stochastic variability. Parameters: *β* = 6 × 10^−4^ and *γ* = 3 × 10^−4^.

**TABLE I.**
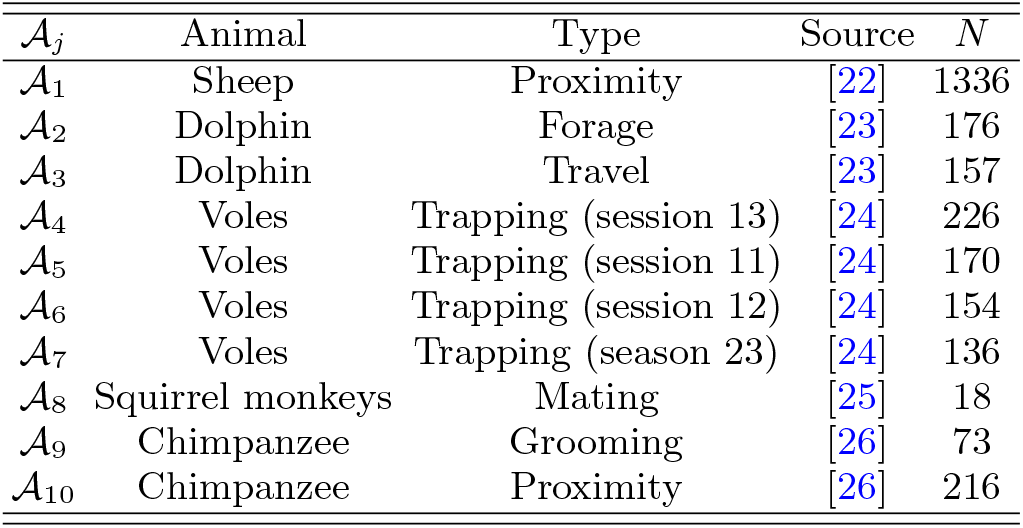
Animal social networks A_j_ used in FIG. 6. Each network exhibits degree heterogeneity and contains super-spreaders. Here, *N* denotes the size of the largest connected component. The ten networks were selected from the database in Ref. [13] as those with the largest values of *Z*_A_, excluding the star graphs.

Increasing infectivity causes the ratio *Z* to decrease and approach *Z*_A_, strengthening the suppression effect. As a result, the average fixation probability curve *ρ*(*s*) becomes flatter due to the stronger suppression. In other words, higher *R*_0_ results in weaker selection, making it harder for advantageous strains to sweep and fix, and enhancing the presence of disadvantageous strains. In fact, we have shown that in the star graph, as *R*_0_ increases leading to an increase in the expected size of the infected population, the suppression factor scales as *Z* ∼ 1*/N*_i_.

The decrease of the suppression ratio with increasing infectivity can be understood physically as follows. Consider a heterogeneous network with only a few super-spreaders. When infectivity is small, infection is mostly sustained on the super-spreaders, while low-degree nodes are infected only rarely. Thus, the average infected sub-population is composed mainly of high-degree nodes and has relatively little degree heterogeneity. As a result, the suppression of selection is weak. As infectivity increases, non-super-spreader nodes also begin to participate more frequently in the infected subpopulation. The infected subpopulation then contains both high-degree and low-degree nodes, making it more heterogeneous. This increased degree heterogeneity strengthens the suppression of selection.

## Appendix D Real world networks

### 1. Suppression of selection in empirical networks

The animal social network repository [13] contains 1213 networks. FIG. 2 shows the distribution of *Z*_A_ across all of these networks. Since we have argued that *Z*_A_ directly affects *Z*, suppression in *Z*_A_ provides a useful structural indicator of selection suppression. However, beyond this structural prediction, we also need to verify the corresponding values of *Z* on these networks using fixation simulations. To do this, we performed fixation simulations while varying the selection strength to estimate *Z*_s_, as described in Appendix B 4, for a subset of animal networks, see TABLE I.

To select the networks presented in TABLE I, for each network in the repository [13], we extracted the largest connected component and computed its size, *N*. We then excluded very small networks with *N* ≤ 15. Networks with a star structure were also excluded, since such networks are expected to produce strong suppression of selection. Next, we arranged the remaining networks in ascending order of *Z*_A_ and selected the top ten networks for further analysis. Then for the selected set of networks we computed the average fixation probability of a mutant infection under different selection strengths. From these fixation probabilities, we obtained the suppression ratio *Z*_s_ as the slope of the fixation probability curve *ρ*(*s*) (averaged over initial mutant locations, see Appendix B 5) and compared it with theoretical value *Z*_t_. As shown in FIG. 6, for most networks *Z*_s_ ≈ *Z*_t_, confirming the accuracy of our theory across diverse animal networks. The level of suppression varies across networks, both in theory and in simulation results. This is because the networks differ in their degree heterogeneity, with some containing a higher abundance of superspreaders than others. We further note that in three of the ten networks, our theoretical approximation deviates significantly from the simulation. In these networks, the steady-state dynamics are characterized by significantly larger fluctuations under the same parameter regime, leading to a breakdown of the approximation.

### 2. Limitations of the theory

Our theory assumes that once a mutant enters the population and ultimately replaces the resident strain within the infected subpopulation, it does not significantly perturb the steady-state infected population size, *N*_i_. To quantify the effective drift and diffusion, we employ a mean-field approximation, which further assumes a degree-uncorrelated network and weak selection (i.e., selection strength close to zero). When the network is small, selection is not weak, or *N*_i_ varies appreciably with mutant abundance, the quantitative predictions of the theory are expected to deviate from simulations. Nevertheless, the hallmark of suppression of selection is encoded in the behavior near *s* = 0, precisely the region where our approximation is most accurate. In this limit, the theory identifies the key quantity controlling suppression, *Z*, and links it to network structure through *Z*_A_. This connection provides a practical guide for anticipating the magnitude of suppression expected for a given network and for specified wild-type infection and recovery rates.

The steady-state behavior exhibits noticeable variability across different networks. While most networks display relatively stable infection levels with small fluctuations around the mean, a subset shows significantly enhanced variability. This suggests that, even under identical parameters, the underlying network structure can strongly influence the magnitude of stochastic fluctuations. In particular, certain networks act as outliers, where the system experiences large deviations from the average behavior (see FIG. 2(b)). We find that *A*_9_ and *A*_8_ are clear outliers: in the former, the network structure promotes enhanced fluctuations, while in the latter, the effect is primarily due to the small size of the host network; notably, *A*_8_ is the smallest network and *A*_9_ is the second smallest among those considered in the main text.

We acknowledge that the networks in the aforementioned repository [13] represent different types of social interactions collected across many independent studies, such as grooming, mating, and proximity. Moreover, the connection weights were defined differently across studies to best represent the specific type of interaction being modeled. Each type of interaction may serve as an appropriate transmission network for some pathogens, but not necessarily for others. For example, an HIV-like transmission network would be more closely approximated by a mating network, whereas a proximity network may not be an appropriate representation. In contrast, for COVID-like respiratory infections, a proximity network, which may also include contacts captured by the mating network, can provide a reasonable approximation of the transmission network. Therefore, when considering a specific pathogen, the relevant type of social interaction should ideally be selected based on the biological mechanism of transmission. However, instead of conducting such pathogen-specific investigations, we ask a more general question. For each type of social network, we consider a hypothetical pathogen whose transmission is well approximated by that network, and then ask whether the corresponding network structure suppresses selection.

## Appendix E Selection–mutation balance

FIG. 8 demonstrates that, even under identical *R*_0_ and same infected subpopulation size *N*_i_ (FIG. 8.(a)), network topology leads to markedly different mutation–selection balance (FIG. 8.(b,c)). In FIG. 8(b), we compare a scale-free network with a fully connected network and observe clear deviations in the stationary mutant distribution. This difference is consistent with the suppression of selection on heterogeneous networks: effective strength of (negative) selection is reduced, allowing mutants to persist at higher abundance. Mean field theory predicts the expected number of mutants on a scale-free network, ⟨*M* ⟩_SF_, to be 1*/Z*_*t*_ times the mean mutant number on fully connected network, ⟨*M*⟩_SF_. This prediction is shown by a dashed black line in panel (b); we observe that in reality, the selection-mutation balance on a scale free network is even higher (see the blue line).

**FIG. 8.**
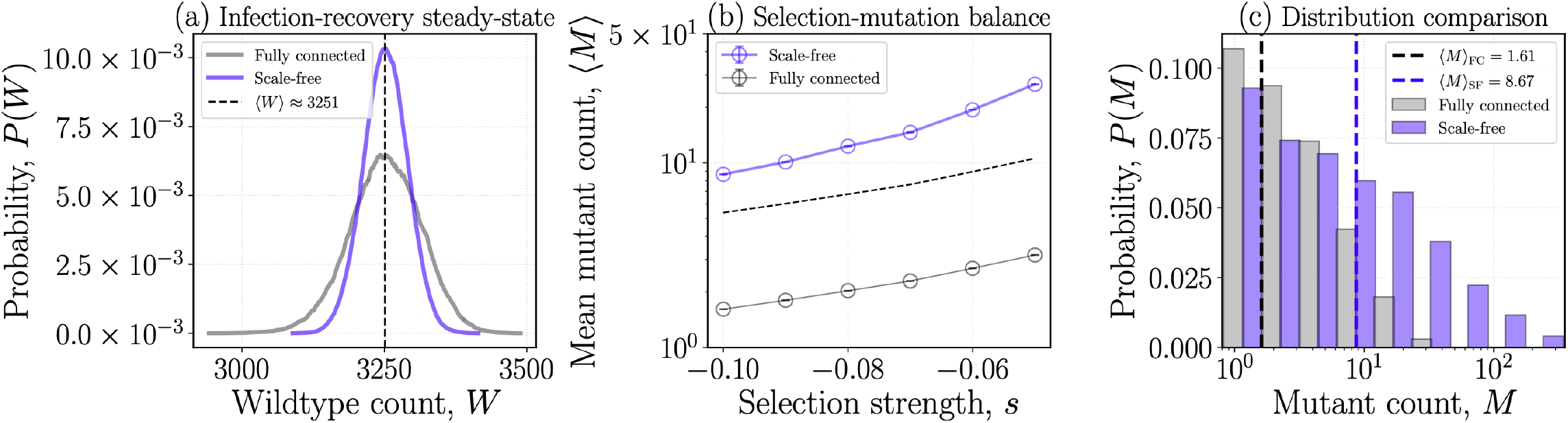
Infection-recovery steady state and selection mutation balance. We consider a scale-free network of size *N* = 5 × 10^3^ and a fully connected network of size *N* = 7.253 × 10^3^. The recovery rate is fixed at *γ* = 3 × 10^−4^ for both networks. To maintain the same wild-type infected subpopulation size *N*_i_ = 3.251 × 10^3^ and basic reproduction number *R*_0_ = 1.81, the infection rate is set to *β* = 6 × 10^−4^ for the scale-free network and *β* ≈ 7.5 × 10^−8^ for the fully connected network (see Appendix A 3). Panel (a) shows the distributions of the wild-type counts for both networks. The distributions are obtained from 2 × 10^7^ time steps after the system has reached the (only wildtype) infection–recovery steady state. We observe that the mean wild-type count, ⟨*W*⟩, attains the same value in both networks. Panel (b), in contrast to panel (a), incorporates disadvantageous mutations and shows the average count of disadvantageous mutants, ⟨*M*⟩, as a function of the selective disadvantage *s*. Mutants are generated from the wild type at a rate *µ* = 5 × 10^−5^. The vertical error bars represent standard errors, which are smaller than the marker size. The dashed black curve is obtained by multiplying the gray curve by 1*/Z*, where *Z* ≈ 0.30 is the suppression factor of selection computed for the scale-free network. Panel (c) shows a representative comparison of the mutant count distributions for a selective disadvantage *s* = −0.1 on both networks. In panels (b, c), − 14.4 × 10^9^ data points were used for any single parameters’ choice.

In the context of the birth–death process, it has been shown [27] that amplification or suppression of selection, defined via fixation probabilities, does not necessarily translate to mutation–selection balance. The infection process considered here is more complex than the standard birth–death setting due to the presence of a fluctuating population size, accounting for a systematic deviation from the mean field theory that is observed.

